# Recurrent pre-leukemic deletions in myeloid malignancies are the result of DNA double-strand breaks followed by microhomology-mediated end joining

**DOI:** 10.1101/2020.01.01.888610

**Authors:** Tzah Feldman, Akhiad Bercovich, Yoni Moskovitz, Noa Chapal-Ilani, Amanda Mitchell, Jessie JF Medeiros, Nathali Kaushansky, Tamir Biezuner, Mark D Minden, Vikas Gupta, Amos Tanay, Liran I Shlush

## Abstract

The mechanisms underlying myeloid malignancies deletions are not well understood, nor is it clear why specific genomic hotspots are predisposed to particular deletions. In the current study we inspected the genomic regions around recurrent deletions in myeloid malignancies, and identified microhomology-mediated end-joining (MMEJ) signatures in recurrent deletions in *CALR, ASXL1* and *SRSF2 loci*. Since MMEJ deletions are the result of DNA double-strand breaks (DSBs), we introduced CRISPR Cas9 DSBs into exon 12 of ASXL1, successfully generating recurrent *ASXL1* deletion in human hematopoietic stem and progenitor cells (HSPCs). A systematic search of COSMIC dataset for MMEJ deletions in all cancers revealed that recurrent deletions enrich myeloid malignancies. Despite this myeloid predominance, we provide evidence that MMEJ deletions occur in multipotent HSCs. An analysis of DNA repair pathway gene expression in single human adult bone marrow HSPCs could not identify a subpopulation of multipotent HSPCs with increased MMEJ expression, however exposed differences between myeloid and lymphoid biased progenitors. Our data indicate an association between MMEJ-repaired DSBs and recurrent MMEJ deletions in human HSCs and in myeloid leukemia. A better understanding of the source of these DSBs and the regulation of the HSC MMEJ repair pathway might aid with preventing recurrent deletions in human pre-leukemia.

---

Human aged HSPCs are prone to clonal expansion due to the acquisition of recurrent mutations. This phenomenon is known as age related clonal hematopoiesis (ARCH)^1-3^. Somatic pre-leukemic mutations (pLMs) do not usually spread randomly across the possible physical positions of a gene, but rather occur at apparent mutational hotspots. Whether mutational hotspots are the result of a specific selective advantage, or are due to increased mutation rate in specific positions remains unclear for most pLMs. The majority of pLMs are nonsynonymous single nucleotide variants (SNVs)^4,5^, however other pLMs are due to recurrent deletions. While the mechanistic explanation for SNVs in cancer has been studied^6^, the mechanisms leading to recurrent deletions in myeloid malignancies are less characterized. The current study sought to identify deletion signatures in myeloid malignancies that would shed light on the origins of these recurrent variants. Two main mechanisms for somatic deletions in cancer are polymerase slippage, mainly in repetitive elements (microsatellite (MS) signature)^7^, or by the error prone process of DSB DNA repair^8,9^.

Analysis of deletion signatures of recurrent myeloid deletions (Fig. 1a) revealed that the most common deletions carry an MMEJ repair signature (Fig. 1b). Based on current knowledge, MMEJ deletions are the result of DSBs ^**Error! Bookmark not defined.**^. Accordingly, we hypothesized that introducing DSBs to HSPCs would result in recurrent myeloid deletions. The introduction of DSBs into the *ASXL1* gene using CRISPR Cas9 in primary human CD34+ HSPCs isolated from 65- and 4-year-old donors resulted in high frequencies of canonical MMEJ deletion (defined here as recurrent MMEJ deletion in humans) in *ASXL1* (Fig. 1c). To better understand whether the non-random distribution of deletions in HSPCs was the result of a selective advantage of canonical MMEJ deletion or of a dominant repair mechanism, the experiment was repeated using the K562 cell line followed by single-cell sorting and expansion. Deletion distributions were quantified from both bulk cells (four days after DSBs) and single-cell derived colonies (one month after DSBs). Similar frequencies of the canonical MMEJ deletions were maintained in bulk and single-cell colonies, suggesting that MMEJ repair dominance was the reason for this non-random distribution rather than it having a selective advantage (Extended Data Fig. 1).

**Figure 1.**
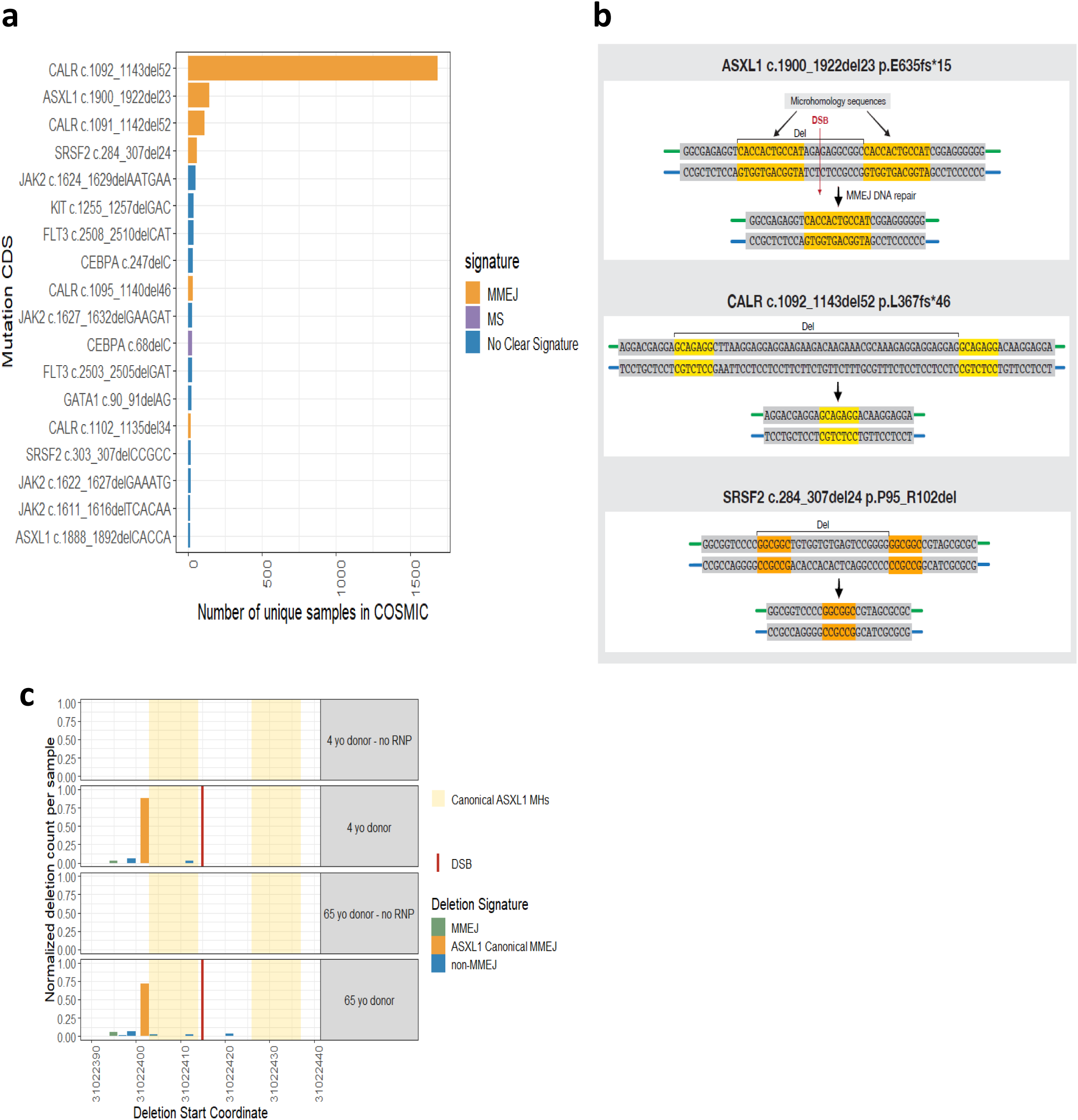
Recurrent deletions in myeloid malignancies share a similar signature of MMEJ mediated DSB repair, successfully recapitulated in primary human CD34+ HSPCs. **a**, Absolute number of samples carrying somatic deletion (represented by gene and mutation CDS (coding DNA sequence) names) in myeloid malignancies identified in 10 or more samples in COSMIC dataset. Deletion signatures are Microhomology Mediated End Joining (MMEJ) (orange), Microsatellites (MS) (purple) and deletions with no clear signatures (blue). **b**, MMEJ deletion signature in *ASXL1, CALR* and *SRSF2* genes. Upon double-strand break (DSB) (red arrow) at genomic loci located between two microhomology sequences (orange and yellow), DNA repair (vertical black arrow) involves a deletion of one microhomology and the sequence between the two microhomology sequences, resulting in canonical MMEJ deletions. **c**, Deep targeted sequencing (read depth 5000X) in primary human CD34+ HSPCs isolated from 65- and 4-year-old donors following CRISPR Cas9 DSBs at specific position (vertical red line) and ‘no RNP’ control samples. Deletion frequency and start genomic positions of the *ASXL1* canonical MMEJ deletion (orange), other MMEJ deletions (green) and deletions with no clear signature (blue) out of the total number of deletions. Canonical ASXL1 Microhomologies (MHs) (orange background) are marked along the *ASXL1* sequence.

While canonical MMEJ deletion in *ASXL1* was most frequently observed in our *in vitro* system, other MMEJ deletions of *ASXL1* never previously reported in humans were also generated, including some truncating variants. It is known that truncating variants across the entire exon 12 of *ASXL1* are pathogenic^10,11^. However, the fact that other MMEJ frameshift deletions do not occur in myeloid malignancies (Fig. 1a) suggests that DSBs at specific positions are responsible for the high prevalence of canonical MMEJ deletions. We therefore designed sequential DSBs along the hotspot regions of the *CALR, ASXL1* and *SRSF2* genes. In both *ASXL1* and *SRSF2*, DSBs located near the midpoint between the two microhomology (MH) sequences produced the highest frequencies of canonical deletions (Fig. 2a, b). We were unable to recapitulate the *CALR* canonical deletion *in vitro* while other MMEJ deletions were generated, probably because guides could be designed at the midpoint between the MHs (Fig. 2c).

**Figure 2.**
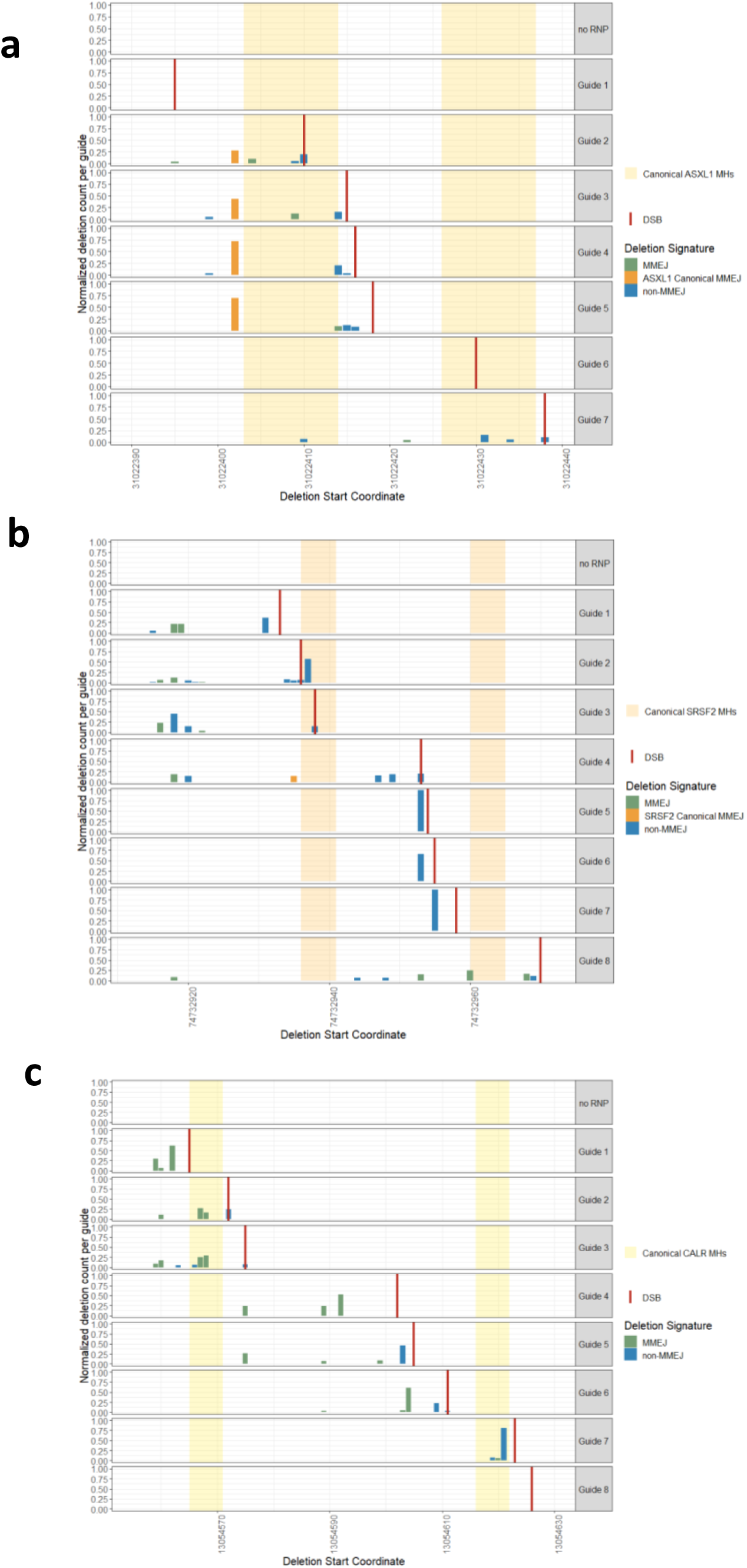
Deletion distribution in *ASXL1* and *SRSF2* genes following sequential CRISPR Cas9 DSBs in K-562. **a b c**, Deletion frequency and start genomic positions of *ASXL1* (a) *SRSF2* (b) and *CALR* (c) canonical MMEJ deletions (orange), other MMEJ deletions (green) and deletions with no clear signature (blue) out of the total deletions assessed by deep targeted sequencing (read depth 5000X) in the K562 cell line following sequential CRISPR Cas9 DSBs (vertical red lines) along *ASXL1* (a) and *SRSF2* (b) and *CALR* (c) genes and ‘no RNP’ controls. Canonical Microhomologies (MHs) (orange and yellow backgrounds) are marked along the sequences.

The specificity in the location of the DSBs raises the possibility that they might occur due to exposure to chemotherapy, like secondary acute myeloid leukemia (AML) which is driven by specific MLL translocations on chromosome 11 after exposure to topoisomerase inhibitors^12^. To test this hypothesis, we analyzed sequencing data from 5,649 cancer patients reported in *Coombs et al.*^*13*^. Canonical MMEJ deletions in *CALR, SRSF2* and *ASXL1* were identified in 16 of the 5,649 patients (*ASXL1* c.1900_1922del23 N=11, *CALR* c.1092_1143del52 N=4, *SRSF2* c.284_307del24 N=1) for 11 of which clinical data was available. In the MMEJ group, 4/11 received chemotherapy prior to sample acquisition. These ratios were not significantly different when compared to individuals without MMEJ deletions (3,610/5,649). Furthermore, canonical MMEJ deletions in *CALR, ASXL1* and *SRSF2* could not be identified in K562 cells exposed to 20Gy irradiation.

Therefore, while neither chemotherapy nor irradiation enriched MMEJ deletions in our analysis, other sources of exogenous stress cannot be excluded as leading to these mutational signatures^14^. To further explore the origin of DSBs leading to MMEJ signatures in myeloid malignancies, we hypothesized that such signatures might exist in other malignancies, thereby shedding light on the mechanisms of DNA DSBs. To identify recurrent MMEJ deletions in cancer, we analyzed all deletions reported in COSMIC, systematically searching for MMEJ deletion signatures (Fig. 1b) with MHs of at least 5bp, which has been suggested as the shortest MH enabling MMEJ repair^8^. While the majority of deletions in cancer carried an MS signature, only 3,062 (2.5%) samples with an MMEJ signature were identified from a total of 119,756 unique samples reported (Supplementary Table 1). Of the MMEJ deletions, 1,927 (63%) were due to canonical deletions in *CALR ASXL1* and *SRSF2*. Eight COSMIC recurrent MMEJ deletions (defined as variants reported in 10 or more different samples) were identified in six different genes (*CALR, ASXL1, SRSF2, TP53, TSC2* and *ARID1A*) without a clear dominance for any specific cancer tissue other than myeloid malignancies (Fig. 3a, Supplementary Table 1). Myeloid recurrent MMEJ deletions were reported in more samples than non-myeloid MMEJ deletions. However, since this might be due to differences in the amount of sequencing data available for different genes in COSMIC, we calculated the proportions of MMEJ deletions and non-MMEJ deletions per gene and discriminated between myeloid genes with a significantly higher MMEJ dominance –such as *CALR* (87%) *SRSF2* (63%) and *ASXL1* (34%) – and TP53, the gene with the highest number of non-myeloid samples with an MMEJ signature (2.4%, p. value <0.0001 for each gene) (Fig. 3b).

**Figure 3.**
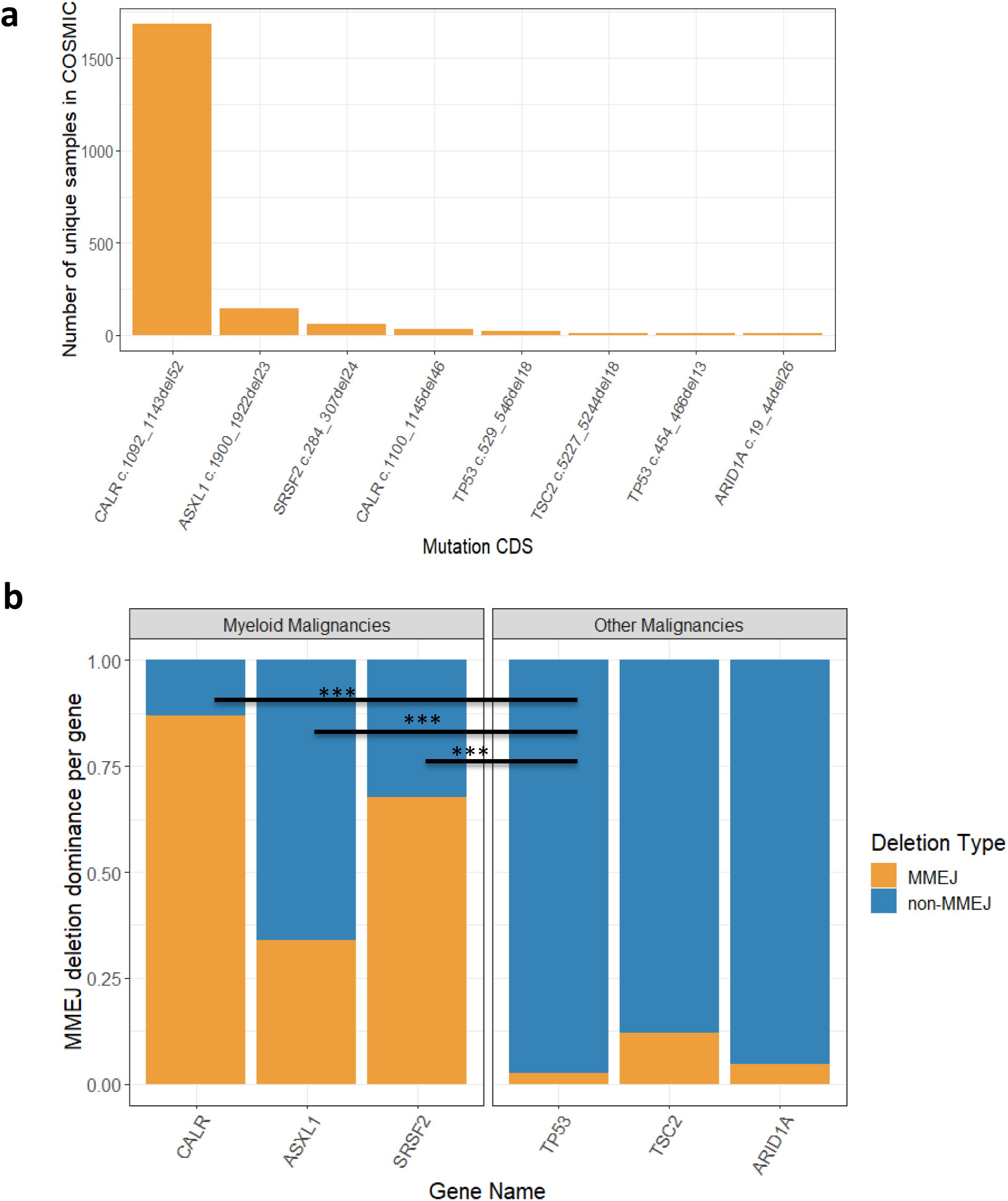
Recurrent MMEJ deletions occur predominantly in myeloid malignancies. **a**, Absolute number of samples carrying deletions with MMEJ signature (represented by gene and mutation CDS (coding DNA sequence) names) identified in 10 or more samples in COSMIC dataset in all cancers. **b**, The proportion of deletions with MMEJ signatures (orange) and non-MMEJ signature (blue) out of the total number of deletions reported for each gene presented in **a.** Significant proportion differences (chi square analysis) are defined as p<0.0005 and are indicated by asterisks (***).

Based on these findings we concluded that the *CALR, ASXL1* and *SRSF2* deletions share an MMEJ signature rarely identified in other cancer tissues. We hypothesized that this myeloid predominance might be related to a specific, shared myeloid-biased cell of origin. We analyzed published sequencing data from healthy individuals and identified the canonical MMEJ deletion in *ASXL1* occurring in three of 124 pre-AML cases 10.7, 8.8 and 1.7 years prior to AML diagnoses (Supplementary Table 2) and none among the 676 controls^4^. These data suggest that long-lived HSCs are the cell of origin for canonical MMEJ deletion in *ASXL1*. To further validate these findings, we analyzed variant allele frequencies (VAFs) of the canonical MMEJ *CALR* deletion in isolated HSPCs and mature cells from two different myelofibrosis (MF) donors. *CALR* deletion was identified among HSCs, more committed progenitors, and mature myeloid and lymphoid cells (Fig. 4a). We transplanted CD34 positive cells from one of the donors into NOD/SCID/IL-2Rgc-null (NSG) mice and after 16 weeks observed a multi-lineage graft. Taken together, these data undermined our hypothesis regarding the cell of origin since the presence of both *ASXL1* and *CALR* MMEJ deletions in multipotent HSCs cannot explain the myeloid predominance of MMEJ deletions.

**Figure 4.**
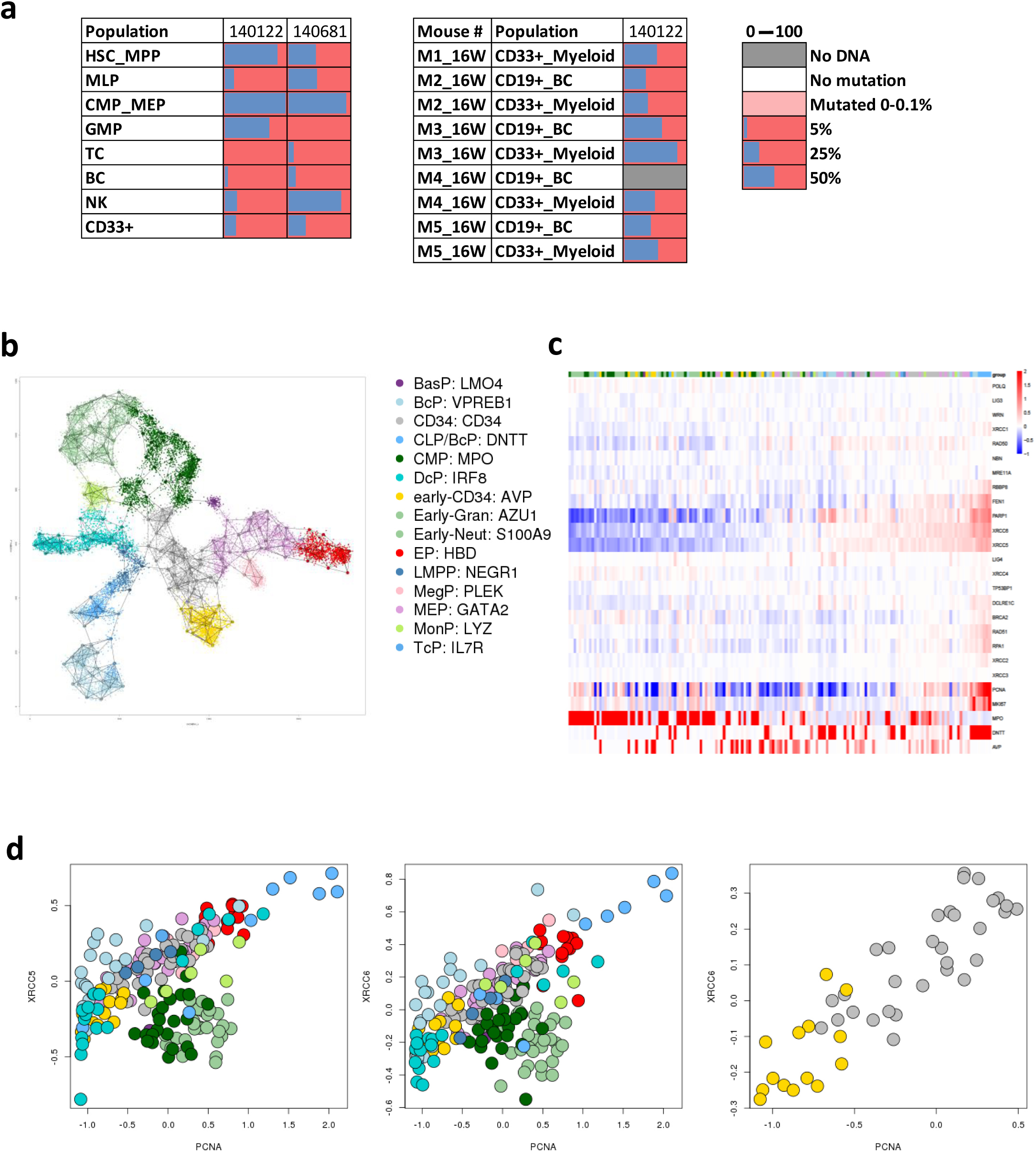
MMEJ canonical deletions are detected in multipotent HSCs. **a**, Variant allele frequency (VAFs) (%) of the canonical *CALR* MMEJ deletion as detected by droplet digital PCR (ddPCR) in various cell populations sorted directly from peripheral blood of two myelofibrosis (MF) patients (samples 140122 and 140681), or in myeloid (CD33+) and B-cells (BC, CD19+) isolated from xenografts generated in NGS mice 16 weeks post intrafemoral transplantation of sample 140122. HSC/MPP, haematopoietic stem cell/multipotent progenitor; MLP, multi-lymphoid progenitor; CMP/MEP, common myeloid progenitor/megakaryocyte erythroid progenitor; GMP, granulocyte monocyte progenitor; TC, T cell; BC, B cell; NK, Natural killer cell. Black squares indicate populations without sufficient DNA amounts for variant detection; dark grey bars indicate VAF. **b**, 2D projection of annotated metacells after filtering for HSPCs from the full HCA immune census dataset. Markers used for annotation are shown to the right. BasP, basophil progenitor; BcP, B-cell progenitor; CLP, common lymphoid progenitor; CMP, common myeloid progenitor; DcP, dendritic cell progenitor; Early-Gran; early granulocyte; Early-Neut, early neutrophil; EP, erythroid progenitor; LMPP, lymphoid-primed multipotential progenitors; MegP, megakaryocyte progenitor; MEP, megakaryocyte erythrocyte progenitor; MonP, monocyte progenitor; TcP, T-cell progenitor **c**, Log fold change (lfp) in UMI content for DSBs repair genes in the various HSPCs metacells. Genes markers of proliferation (PCNA and MKI67) and gene markers (MPO, DNTT, AVP) lfp’s are also shown. **d**, Scatter plots of HSPCs metacells’ lfp values for XRCC5 and XRCC6 vs. proliferation marker, with XRCC6 also shown for the less differentiated progenitors.

Since our results support HSCs as MMEJ deletions’ cell of origin, we hypothesized that specific subpopulations of HSPCs are capable of MMEJ repair due to differential expression levels of DNA repair pathways. While previous studies on DNA repair in rodent^15^ and human^16^ focused on bulk HSPCs, we aimed to characterize the DNA repair landscape in human BM at the single cell level. To address this issue we analyzed the Human Cell Atlas Consortium’s immune census dataset (https://preview.data.humancellatlas.org/), which consists of roughly 310,000 single-cell RNA profiles from adult human BM (Extended Data Fig. 2). Using the Metacell pipeline^17^, we isolated and annotated (Fig. 4b, Extended Data Fig. 2) putative HSPCs (CD34+ N=19293) from BM. Analyzing differences in relative gene expression of key factors of the DNA repair pathways^18^ together with markers of proliferation (PCNA and MKI67) (Fig. 4c), we identified heterogeneity in the expression levels of canonical non-homologous end joining (c-NHEJ) repair pathway (mainly in *XRCC5, XRCC6*) across HSPCs. In early HSCs, expression levels of all repair pathways correlated with replication/cell cycle. MMEJ pathway expression is relatively homogenous across HSPCs with less clear association with cell cycle. On the other hand, differences in expression levels of c-NHEJ factors were observed between lymphoid-biased progenitors and myeloid-biased progenitors (Fig. 4c). While the c-NHEJ expression is increased with proliferation among B-cell progenitors such phenomenon was not observed among myeloid progenitors (Fig. 4d). These differences can be explained by the dependency of B /T cell receptor rearrangement on c-NHEJ repair during lymphoid cells development^19^. Since MMEJ repair was shown to be active when the c-NHEJ pathway is knocked out^20,21^, it is possible that HSPCs with low c-NHEJ expression could be primed for MMEJ repair. However, these results could not identify the exact cell population at risk for MMEJ error-prone repair.

In conclusion, we found that the three common pre-leukemic somatic deletions in human myeloid malignancies *ASXL1* c.1900_1922del23, *SRSF2* c.284_307del24 and *CALR* c.1092_1143del52 share a similar deletion signature (termed here canonical MMEJ deletions). In a CRISPR model of DNA DSBs, canonical MMEJ deletions in *ASXL1* and *SRSF2* were generated following DSBs in specific genomic loci. Our study also provide evidence that MMEJ deletions in *ASXL1* and *CALR* originate in early HSCs. We further identified heterogeneity in expression levels of DNA repair pathways, mainly in c-NHEJ genes across single HSPCs. Our findings raise questions which merit further research: 1) Why do DSBs occur to begin with? 2) Why are DSBs repaired in HSCs by the MMEJ error-prone pathway? 3) Why are MMEJ canonical deletions enriched in age-dependent myeloid malignancies? 4) Are other deletions and structural variants in myeloid malignancies the result of MMEJ or other repair pathways?

As the canonical MMEJ deletions were not produced by radiotherapy *in vitro* and were not enriched among individuals exposed to chemotherapy, it is possible that other DSB sources – including, replication fork stalling during DNA replication – might explain these deletions^**Error! Bookmark not defined.**,22,23^. Studies on mouse aged HSPCs suggested that they carry more DSBs and Gamma H2AX foci due to altered dynamics of DNA replication forks^24^. Interestingly, polymerase theta (coded by the POLQ gene), a key factor in the MMEJ pathway, was shown to protect against DNA rearrangements during replication at the expense of small deletions at G4 DNA sites^25^. Other DSB sources may be chromatin conformations, enhanced transcription, RNA:DNA hybrid structures, excision of transposable elements and endonuclease mediated programmed DSBs ^**Error! Bookmark not defined.**^. These mechanisms should be further studied in the context of ageing human HSCs and the accumulation of DSBs in specific hotspots.

An analysis of COSMIC data revealed an enrichment of recurrent MMEJ deletions in myeloid malignancies. This myeloid enrichment could be due to either a specific HSC subpopulation prone to MMEJ repair or to high probability of DSBs occurrence in a subset of HSCs at specific positions. While we could not detect a specific HSPC subpopulation prone to MMEJ repair, possibly due to the scarcity of such population, future studies should further explore the significance of the MMEJ pathway in early HSCs in the context of human pre-AML. It remains unclear, however, why canonical MMEJ deletions are enriched in adult leukemia (but not in pediatric leukemia) and mainly in three specific hotspots and not in other pre-leukemic genes enriched with deletions (*DNMT3a, TET2*). One limitation of our study was the reliance on COSMIC dataset which is not necessarily representative. Future studies should therefore focus on novel methods to identify MMEJ deletions from whole genome and whole exome sequencing data, and to further expand the search to copy number variation^26^ and translocations in myeloid malignancies. Another limitation was the use of CRISPR Cas9 as a model for DSBs, as CRISPR Cas9-mediated DSBs probably differ from DSBs in human cells. Recent studies demonstrated that genome-wide CRISPR editing enriches MMEJ deletions^27^ while human somatic deletions do not (Supplementary Table 1). Consequently, future studies should identify other techniques (besides CRISPR and irradiation) to initiate DSBs and study their effects.

Our findings support the growing evidence that cancer mutations do not occur randomly and that their physical position is determined not just by the selective advantage they provide. Understanding the mechanisms of early cancer mutations is crucial for cancer prevention. Discovering the reason DSBs occur in specific hotspots and why they are repaired by the deleterious MMEJ pathway could potentially prevent MMEJ-derived myeloid malignancies.

## Supplementary information

Supplementary Table 1 – Clinical summary and deletion signatures of 119,756 deletions reported in COSMIC dataset.

Supplementary Table 2 – Clinical and genetic characteristics of pre-AML samples carrying *ASXL1* canonical MMEJ deletion.

## Acknowledgements

The authors wish to thank Prof. John Dick and Dr. Ayal Hendel for fruitful discussion and support. Biological samples from Mobilized PBSC autologous transplant products were generously provided by Dr. Mark Minden. L.S. is the incumbent of The Ruth and Louis Leland career development chair.

This research was supported by the EU horizon 2020 grant project MAMLE ID: 714731 and LLS Grant ID: RTF6005-19.

## Author contribution

T.F designed and developed the study, performed CRISPR experiments, cells culture and maintenance, deep targeted sequencing, analyzed sequencing data, performed variant calling, analyzed COSMIC data and wrote the manuscript. A.B performed single-cell RNA analysis. Y.M performed CRISPR experiments, cells culture and maintenance, deep targeted sequencing. N.I.C provided bioinformatics support and wrote the matlab code for MMEJ detection. N.K revised the paper and contributed to data interpretation. T.B provided sequencing and technical support. A.M and J.M Performed xenotransplantation experiments, cell sorting and ddPCR. M.D.M and G.V enabled sample acquisition A.T. supervised single-cell RNA analysis L.I.S. designed and supervised the study and wrote the manuscript.

## Methods

### 1. Samples

Biological samples from Myelofibrosis (MF) patients were collected with informed consent according to procedures approved by the Research Ethics Board of the University Health Network (REB 01-0573-C). Mobilized peripheral blood autologous transplant products were collected with informed consent according to procedures approved by the University health network ethics committee protocol # 15-9633, and Weizmann institute of science IRB protocol #337-1.

### 2. CRISPR experiments

Targeted genome editing in K562 cells was performed by using CRISPR Cas9 system according to protocols described elsewhere^28,29^ with slight modifications.

#### 2.1 Guides design and plasmid preparations for K562 experiments

20bp sgRNA sequences were designed around the genomic loci of interest using DESKGEN algorithm (https://www.deskgen.com/landing/#/login) (Extended Data Table 1). Sense and antisense oligonucleotides for each sgRNA with overhangs compatible to Bbsi-digested px330 were designed and ordered from IDT.

Each oligos pair was further phosphorylated and annealed using T4 PNK (NEB) and T4 Ligation Buffer (NEB). Phosphorylation and annealing reaction was performed at 37°C for 30 min, followed by 95°C for 5 min and ramping down to 25°C at 5°C /min. Annealed oligo pairs were then ligated into a previously Bbsi digested px330 plasmid. Per reaction, 50ng digested px330 was mixed with 1:250 diluted oligo duplex with 2X quick ligation buffer (NEB) and quick ligase (NEB) at 16°C overnight.

BioSuper DH5α competent cells (Biolab) were transformed with Px330. Bacteria was resuspended and plated on LB agar AMP dishes and incubated at 37°C over-night. Colonies were then screened and grew in 2-3 ml LB + Ampicillin at 37°C overnight in a shaker (250rpm). For each colony (guide), plasmid DNA was extracted using the QIAprep Spin Miniprep standard protocol (Qiagen, cat. No. 27104). To validate the presence of the desired inserts, Sanger sequencing reactions were performed for each plasmid using the U6 promoter primer ACTATCATATGCTTACCGTAAC.

#### 2.2 Electroporation reactions

All electroporation reactions were performed using the 16-strip Lonza 4D nucleofector kit. Electroporation reactions on primary CD34+ enriched and K562 cells were performed by using a synthetic *ASXL1* targeted sgRNA ordered from IDT and resuspended in IDTE buffer to a final concentration of 100uM. K562 sequential sgRNA experiments, were done using purified px330 plasmids at 2ug/reaction. For each unique sample per electroporation, a control sample containing the same cell amount with no RNP/plasmid, underwent electroporation at the same conditions was sequenced.

#### 2.3 K562 cell line experiments

K562 cell line experiments using px330 plasmids, were performed according to the manufacture recommendations. Briefly, K562 cells were cultured in ATCC-formulated Iscove’s Modified Dulbecco’s Medium containing 10%FBS and 1% pen-strep and split to 300,000 cells/ml 48h prior to electroporation. On electroporation day, cells were counted and resuspended in SF nucleofector solution. 200,000 cells /reaction were resuspended in 20ul SF solution and mixed with 2ug px330 plasmids.

K562 cell line experiments involving synthetic sgRNA targeted to *ASXL1* were done according to IDT recommendations. *ASXL1* synthetic sgRNA was ordered from IDT according to the following sequence: AGGTCACCACTGCCATAGAG. RNP complexes were generated by mixing 2.1ul PBS, 1.2ul 100uM sgRNA and 1.7ul synthetic CAS9 (IDT) per reaction, followed by incubation at room temp for 10-20 min. Complexed RNPs were then transformed to 4°C. 5ul of complexed RNP were mixed with 20ul SF solution containing 200,000 K562 cells Per reaction as was described.

Program FF-120 was used. Following electroporation, cells were washed and cultured. 4 days following electroporation, cells lysate was extracted by mixing each sample pellet with 30ul of 50mM NaOH and heating at 99°C for 10 min. Then, cells lysate was cooled on ice and 2ul 1M Tris at ph=8 was added to each reaction. All lysates were next served as a template for library preparations.

#### 2.4 K562 single cell sorting and expansion

Bulk electroporated K562 cells underwent sorting for live single cells using BD FACS Aria III sorter. Sorted cells were plated onto two 96-well plates in 200ul/well culture medium (described in 2.3). Cells maintained by replacing 100ul medium from each well once a week. Once enough cells were observed, lysate was extracted.

#### 2.5 Primary CD34+ enriched cells electroporation

CD34+ enrichment and electroporation was done according to IDT protocol “Electroporation of primary human CD34+ hematopoietic stem and progenitor cells” contributed by Dr. Ayal Hendel. 48 hours prior to electroporation, CD34+ cells were isolated from mononuclear cells derived from mobilized PBSC autologous transplant products by using CD34 Miltenyi Biotec MicroBead kit. Cell were re-plated in cytokine rich medium in a 24-well plate to reach a density of 250,000cells/ml. CD34+ cells cytokine rich medium was based on SFEMII supplemented with SCF (100 ng/mL), TPO (100 ng/mL), Flt3-Ligand (100 ng/mL), IL-6 (100 ng/mL), streptomycin (20 mg/mL), and penicillin (20 unit/mL). Cells were incubated in a low-oxygen, tissue culture incubator (37°C, 5% CO2, 5% O2) according to IDT protocol. 24h prior to electroporation, cells were counted and cell density was maintained below 10^6 cells/ml. On electroporation day, cells were washed with PBS and counted. 250,000 CD34+ cells per reaction were resuspended in 20ul P3 nucleofector solution and mixed with 5ul RNP complexes that were generated as described under section 1.3. Program DZ-100 was used. Following electroporation, cells were washed with 80ul pre-warmed cytokine rich medium and transferred into a U-bottom 96-well plate containing 100ul pre-warmed cytokine rich medium. CD34+ cells were cultured for 2 additional days in a low-oxygen incubator followed by a lysate generation that was subsequent to library preparations.

### 3. K562 irradiation experiment

K562 cells were cultured in 96-well plates in 200ul medium/reaction. Cells in four different wells were irradiated with gradient dosages of 0, 5, 10 and 20Gy. Cells were cultured for additional four days. Then, lysate was extracted for targeted sequencing.

### 4. Targeted Sequencing

Dual indexed illumina Libraries were generated using two-step PCR procedure. 1^st^ PCR primer prefix sequences and 2^nd^ PCR primer sequences were, as previously described^30^ with a modified protocol. In short, target-specific primers were designed by Primer3plus (http://www.bioinformatics.nl/cgi-bin/primer3plus/primer3plus.cgi) and were ordered with the described 5’ prefixes^30^ (IDT) (Extended Data Table 1). 1^st^ PCR was applied to target the regions of interest. The reaction mixture was composed of a PCR ready mix (using NEBNext^®^ Ultra™ II Q5^®^ Master Mix, NEB), lysate product and a final primer concentration of 1uM each. PCR protocol was as follows: 98°C for 30 sec, followed by 40 amplification cycles of 98°C for 10 sec,65°C for 30 sec and a final elongation at 65°C for 5 min. Following dilution of the 1^st^ PCR products with nuclease free water (1:1000), a 2^nd^ PCR was performed using primers composed of Illumina sequencing primers, indexes and adapters, under the same conditions as the 1^st^ PCR with the exceptions of final primer concentration of 0.5uM each and 20 cycles of amplifications. Samples were pooled together at equal volume. Pooled library sizes were selected (2% gel, BluePippin, Sage Science) and sent for 2 × 150-bp deep sequencing (Miseq, Illumina).

### 5. Variant Calling

2 × 150-bp pair-end reads deep sequencing data (∼5000X depth) from Illumina platform were converted to fastq format. Minimap2.1 algorithm^31^ was applied for alignment of the processed fastq files to hg19 genome based targeted sequences resulting in sam files that were further sorted and indexed using pysam 0.15.1 (https://github.com/pysam-developers/pysam). All reads from sorted bam files were assigned to new read groups using picard 2.8.3 ‘AddOrReplaceReadGroups’ command (http://broadinstitute.github.io/picard). In order to avoid misalignments, local realignment was preformed using GATK3.7 ‘RealignerTargetCreator’ and ‘IndelRealigner’ commands^32^. Mpileup files were generated by samtools 1.8 followed by SNVs and small indels detection using varscan2.3.9 ‘pileup2cns’ command to generate VCF files containing consensus variant calls^33^. Short indels at positions around polyG sequences (position 20:31022441 in ASXL1 and 17:74732955 in SRSF2) identified in control ‘no RNP’ samples in CRISPR experiments were manually excluded from the analyses. Deletion frequencies from bulk cells were obtained by dividing variant allele frequency by the sum of all deletions allele frequencies per sample. For single cell analysis, a threshold deletion VAF ≥ 20% was set for each single cell derived colony. Frequency of colonies carrying deletions were obtained by dividing colony numbers carrying each variant (at ≥20% VAF) by the total colonies number. MMEJ deletion signature from all CRISPR experiments were defined as MH>3bp and no mismatches were allowed.

### 6. Cell Sorting, xenotransplantation and ddPCR analysis

Fluorescence-activated cell sorting of human stem/progenitor and mature cell populations was performed on mononuclear cells from peripheral blood of two MF patients according to the sorting strategy described in details elsewhere^34^. Animal experiments were performed in accordance to the IACUC of the Weizmann Institute, its relevant guidelines and regulations (11790319-2). Eight- to 12-week-old female NOD/SCID/IL-2Rgc-null (NSG) mice were sublethally irradiated (225 cGy) 24 hours before transplantation. CD34+ cells were enriched by magnetic beads (Miltneyi Inc.) and 50,000 cells were injected into the right femur as previously described^34^. Mice were euthanized 16 weeks following transplantation and human engraftment in the injected right femur and non-injected bone marrow (left femur, tibias) was evaluated by flow cytometry. Subpopulations were sorted as previously described^34^.

ddPCR reaction was performed by using probes designed for *CALR* deletion as described elsewhere^35^. Amplified DNA (2ul from a 1:20 dilution of a 16 h REPLI-g Mini Kit whole-genome amplification, Qiagen) from each sorted population was tested in a 96-well plate in duplicate according to the manufacturer’s protocol. Mutant and wild-type sequences were read using a droplet reader with a two-color fluorescein/HEX fluorescence detector (Bio-Rad). The mutant allele frequency was calculated as the fraction of mutant-positive droplets divided by total droplets containing a target. As previously reported^3^ the minimum detection level was 1:1,000 (0.1%). Variants were considered present if there were at least three dots in the mutant fluorescein channel resulting in VAF > 0.1%.

### 7. Single cell RNA-seq analyses

HSPCs RNA-seq profiles were isolated from the HCA immune census BM data based on CD34 expression. A total of 19757 profiles were isolated from the roughly 310,000 BM profiles from 8 different donors (list of isolated profiles used can be found in the supplementary data). To generate metacells from the profiles, we used the MetaCell package14 with parameters as specified below. Feature genes were selected using the parameter T_vm = 0.08 and minimal total UMIs of 100, while excluding genes correlated to lateral effects such as mitochondrial genes, immunoglobulin genes, high abundance, prefix “RP-” genes, cell cycle, type I Interferon response and stress (list of feature genes and excluded genes in supplementary data). The final feature genes, consisting of 527 genes, were used for the computation of the Metacell balanced similarity graph, with parameters k = 60, n_resamp = 500 and min_mc_size = 20. Outliers threshold of T_lfc = 3.5 was used, with 464 profiles deemed as outliers. Next, we annotated the metacell model using hierarchical clustering of the metacell confusion matrix, supervised analysis of enriched genes and analysis of marker genes (Sup. Figures). The metacells and profiles were projected and plotted in 2D using mc2d_K = 40, mc2d_T_edge = 0.02 with a max degree of 6, and colored using thresholds on metacells log enrichment scores (lfp values) for marker genes chosen from common markers and above annotation process. For studying DSBs repair machineries genes expression patterns in HSPCs, we used a heatmap of metacells lfp values for the known repair genes along with markers of proliferation and specific marker genes. Metacells were ordered based on “XRCC5” and “XRCC6” genes mean lfp values. This analysis was performed on all HSPCs metacells, and for specific chosen subsets such as early progenitors (with markers “CD34” and “AVP”, Sup. Figures).

### 8. Cosmic data analysis

COSMIC mutation data containing data from both targeted and genome wide screens was downloaded from https://cancer.sanger.ac.uk/cosmic/download. Deletion table was generated by filtering for rows containing the letters ‘del’ in ‘Mutation CDS’ column using ‘awk’ command. Additional filtering for rows containing genomic coordinates in column ‘Mutation genome position’ was done. This resulted in a table of 210,442 reported deletions. For each case, coordinates with flanking 20bp from both deletion ends (e.g start-20bp, end+20bp) were generated and later used as an input for bedtools getfasta command to generate Fasta file for all COSMIC deletions using hg19 reference genome.

‘MMEJ signature’ detection was done using an in-house matlab code that analyzed each deletion’s flanking sequences for microhomologies (MHs) according to the ‘MMEJ signatures’ described in Fig. 1b.

In addition, the definition of MMEJ deletions included the following criteria:

1. Deletions for which ‘MMEJ signature’ was detected.
2. Deletion length/ MH length > 1
3. MH lengths ≥ 5bp.

Accordingly, cases in which ‘MMEJ signature’ was detected and Deletion length/ MH length ≤ 1 were defined as MS deletions regardless of their MH lengths.

Cases with short MH lengths in which ‘MMEJ signature’ was detected, deletion length/ MH length > 1 and MH lengths < 5bp were combined with cases in which ‘MMEJ signature’ was not detected and considered deletions with no clear signature.

For each reported deletion, minor allele frequencies (MAF) were identified using Annovar tool (https://github.com/WGLab/doc-ANNOVAR) according to the following datasets: AF, ExAC_ALL, Kaviar_AF, ExAC_nonpsych_ALL and AF_popmax. Common SNPs were defined as variants with MAF of 0.0001 or above in at least one dataset and were filtered out. Deletions with no available data in any of these datasets were further included. This filtering resulted in a table of 157,539 deletions. Duplicates were removed according to the combination of values from columns ‘Gene name’, ‘Sample name’ and ‘Mutation genome position’, which resulted in a final table of 119,756 unique deletions (Supplementary Table 1).

For Fig. 1a, a table of all myeloid deletions was generated by filtering the ‘Primary site’ column of the final table to include ‘haematopoietic and lymphoid’ tissue deletions followed by the exclusion of the letters ‘lymph’ from the ‘Primary histology’ column. Signature types were assessed for deletions reported in 10 or more unique samples and a single nucleotide mismatch was allowed in MMEJ signature (mainly in the case of CALR c.1091_1142del52). Identical deletions with multiple ‘Mutation CDS’ values were combined under a uniform name (for example *ASXL1*_c.1888_1910del23 and c.1900_1922del23 were combined under the name *ASXL1* c.1900_1922del23).

For Fig. 3a, a table of all MMEJ deletions was generated by applying an in house matlab script as described to detect MMEJ deletions. Identical deletions with multiple ‘Mutation CDS’ names were combined under a uniform name.

For Fig. 1a and 3a, all identified deletions were arranged using R software according to reported counts in COSMIC after duplicates removal. Variants that were reported in 10 or more unique samples were further filtered according to the following exclusion criteria:

1. Common SNPs at adjacent genomic loci
2. Deletions at intronic or intergenic regions
3. Deletions that were reported in one single paper

All analyses and main figures 1, 2, 3 were generated using R statistical programming environment.

## Extended Data figures

**Extended Data figure 1. Deletion distribution in *ASXL1* following CRISPR Cas9 DSBs in K-562 bulk and single cells.**

**a**,**b**, Deletions frequency (a) or frequency of colonies carrying deletions (b) and start genomic positions of *ASXL1* canonical MMEJ deletions (orange), other MMEJ deletions (green) and deletions with no clear signature (blue) out of the total deletions (a) or total colony number (b) assessed by deep targeted sequencing (read depth 5000X) in bulk (a) and single cells (b) K562 following CRISPR Cas9 DSBs at specific position (vertical red lines) and ‘no RNP’ control samples (a). Canonical Microhomologies (MHs) (orange backgrounds) are marked along the *ASXL1* sequence.

**Extended Data figure 2. Single cell RNA-seq from all Meta cell model.**

**a**, 2D projection of annotated metacells from the entire after filtering for HSPCs from the full Human Cell Atlas Consortium’s immune census dataset which consists of roughly 310,000 single-cell RNA profiles from adult human BM. **b**, markers defining the different metacells.

**Extended Data figure 3. Single cell RNA-seq from all Meta cell model.**

**a b c**, Scatter plots of HSPCs metacells’ lfp values of the different annotation markers for the less differentiated progenitors (a), megakaryocyte erythrocyte progenitors (b), B-cell progenitor/common lymphoid progenitors (c) and common myeloid progenitor (d) vs. CD34.

**Extended Data figure 4. Single cell RNA-seq from all Meta cell model.**

**a b c**, Log fold change (lfp) in UMI content for DSBs repair genes in myeloid and lymphoid progenitors (a) and early HSPCs (b) metacells. Genes markers of proliferation (PCNA and MKI67) and gene markers (MPO, DNTT, AVP) lfp’s are also shown.

**Extended Data figure 5. Single cell RNA-seq from all Meta cell model.**

**a b**, Scatter plots of HSPCs metacells’ lfp values for POLQ (a) and BRCA2 (b) vs. proliferation marker (PCNA).

